# Interacting evolutionary pressures drive mutation dynamics and health outcomes in aging blood

**DOI:** 10.1101/2020.04.25.058677

**Authors:** Kimberly Skead, Armande Ang Houle, Sagi Abelson, Mawusse Agbessi, Vanessa Bruat, Boxi Lin, David Soave, Liran Shlush, Stephen Wright, John Dick, Quaid Morris, Philip Awadalla

**Affiliations:** Ontario Institute for Cancer Research, Toronto, Ontario, Canada; Department of Molecular Genetics, University of Toronto, Toronto, Ontario, Canada; Vector Institute for Artificial Intelligence, Toronto, Ontario, Canada; Department of Mathematics, Wilfrid Laurier University, Waterloo, Ontario, Canada; Department of Immunology, Weizmann Institute of Science, Rehovot, Israel; Department of Ecology and Evolutionary Biology, University of Toronto, Toronto, Ontario, Canada; Princess Margaret Cancer Centre, Toronto, Ontario, Canada; Computational and Systems Biology Program, Memorial Sloan Kettering Cancer Center, New York, NY, United States; The Terrence Donnelly Centre for Cellular and Biomolecular Research; Dalla Lana School of Public Health, University of Toronto, Toronto, Ontario, Canada

## Abstract

A small population of self-renewing, hematopoietic stem cells continuously reconstitutes our immune system. As we age, these cells, or their pluripotent descendants, accumulate somatic mutations; some of these mutations provide selection advantages and increase in frequency in the peripheral blood cell population. This process of positive selection, deemed age-related clonal hematopoiesis (ARCH), is associated with increased risk for cardiac disease and blood malignancies, like acute myeloid leukemia (AML). However, it remains unclear why some people with ARCH do not progress to AML, even when their blood cells harbor well-known AML driver mutations. Here, we examine whether negative selection can play a role in determining AML progression by modelling the complex interplay of positive and negative selective processes. Using a novel approach combining deep learning and population genetic models, we detect pervasive negative selection in targeted sequence data from the blood of 92 pre-AML individuals and 385 healthy controls. We find that the relative proportion of passenger to driver mutations is critical in determining if the selective advantage conferred to a cell by a known driver mutation is able to overwhelm negative selection acting on passenger mutations and allow clones harbouring disease-predisposing mutations to rise to dominance. We find that a subset of non-driver genes is enriched for mildly damaging mutations in healthy individuals fitting purifying models of evolution suggesting that mutations in these genes might confer a protective role against disease-predisposing clonal expansions. Through exploring non driver-centric models of evolution, we show how different classes of evolution act to shape hematopoietic dynamics and subsequent health outcome which may better inform disease prediction and unveil novel therapeutic targets. We anticipate that our results and modelling techniques can be broadly applied to identify both driver mutations and those mildly damaging passenger mutations, as well as help understand the early evolution of cancer in other cells and tissues.

---

Hematopoiesis proceeds through an extensive differentiation hierarchy rooted in a population of hematopoietic stem cells (HSCs) (1). The HSC pool is estimated to consist of between 10,000 to 200,000 cells (2,3) and is among the most productive and tightly regulated populations in the human body. As individuals age, somatic mutations accumulate in HSCs, or in early blood cell progenitors (3–6). Some mutations confer a proliferative advantage to certain cells, clones, in the hematopoietic hierarchy and result in a disproportionate lineage representation in the mature blood cell pool (3–6). This predicted imbalance is observed with increasing frequency as individuals age and accordingly has been called Age-Related Clonal Hematopoiesis (ARCH) or, alternatively, Clonal Hematopoiesis of Indeterminate Potential (CHIP) (3–7). ARCH has been linked to an increased risk for cancer and various cardiovascular (CVD) conditions including inflammation, atherosclerosis, thrombosis and sudden death (3–7). However, only a small proportion of individuals with ARCH progress to disease, and mechanisms driving the transformation to malignancy remain unclear. With respect to cancer, specifically Acute Myeloid Leukemia (AML), the presence of mutations in known AML driver genes at a high frequency is one of the best predictors of later disease onset. However, these same mutations are observed in healthy individuals who display no signs of hematological malignancy (7).

The evolutionary trajectory of somatic mutations in cellular populations is governed by a combination of deterministic processes, selection, and stochastic neutral processes (genetic drift) (8). Mutational profiles from blood afford us a unique opportunity to study somatic evolutionary processes within, and among, individuals. Each blood sample is a reflection of the population history of the aging HSC pool as well as the derived cell populations. Through high-coverage sequencing, we are able to capture the full spectrum of mutational variation within each blood population, including variants segregating at extremely low frequencies which are likely to be the targets of negative selection and which are typically not captured in low-to moderate-coverage sequencing efforts.

The ability to detect and quantify negative selection would allow us to move beyond the comparison of exclusively adaptive versus neutral models, which are conventionally used to model cancer evolution (9–11) and explore more diverse models that consider negative selection (12,13). For example, the genetic background on which driver mutations arise has not been well characterized and could explain variation in outcomes. Indeed, little attention has been paid to the role of mutations occurring in non-driver genes in shaping disease outcomes (14). For the remainder of this paper, we will refer to mutations accumulating in non-driver genes as “passenger mutations”. Some passenger mutations are damaging, and should they occur within a clone carrying a driver mutation, could impact the rate of expansion of the driver-harboring clones (***Figure 1a***) (15). As the majority of mutations (>99.9%) in cancers are passenger mutations, modeling their appearance and contribution to clonal fitness is critical to understanding mutation rates, cellular evolution and disease progression (16). As such, to fully exploit the predictive potential of ARCH for cancer and CVD outcomes, it is critical that we consider the full range of selective events occurring in the mature blood pool.

**Figure 1.**
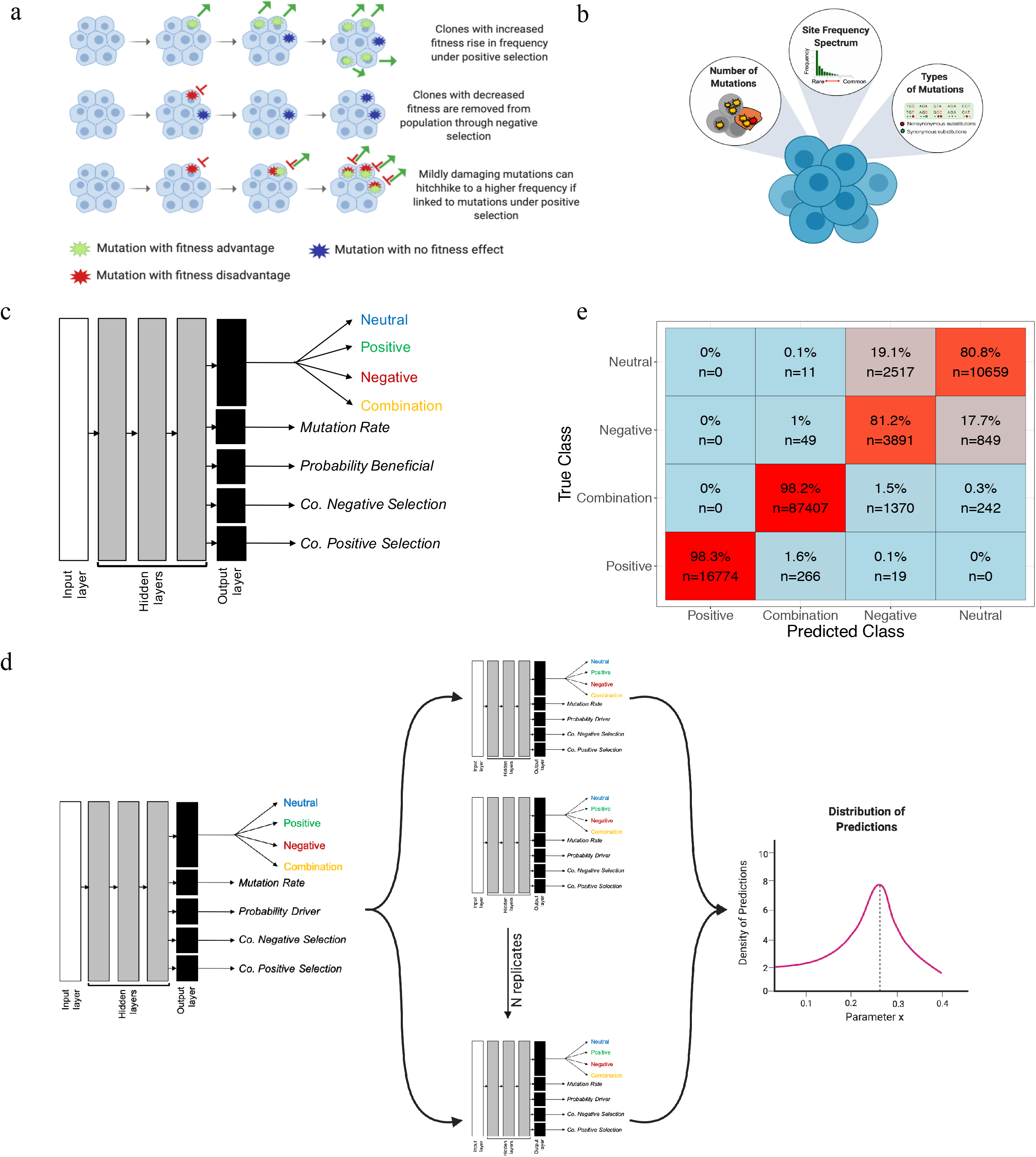
**a)The impact of selection and genetic drift on shaping clonal dynamics.** Cells accumulate somatic mutations in each division. The majority of mutations will be either neutral (blue) or mildly damaging (red). Driver mutations, will increase the fitness of a cell and increase in frequency in the population (green). However, mutations are also able to rise in frequency through genetic drift. **b) Mutation summary statistics extracted from blood cell populations.** Summary statistics fall into three main categories: 1) Counts of mutations in each blood sample, 2) The frequency of mutations (variant allele frequency) and 3) mutation annotation and respective ratios (proportion of missense relative to total missense sites over the proportion of silent mutation relative to total silent sites). A total of 16 summary statistics are extracted from each population. **c) Deep Neural Network Architecture.** Each DNN was trained as a multi-task neural network and classifies a population into one of four overarching evolutionary classes, and predicts four continuous parameters. Each DNN had three hidden layers comprised of 512 nodes each. **d) DNN Ensemble**. We trained a total of ten deep neural networks (DNNs) independently, yet with identical architecture. Through employing an ensemble based approach, we are able to obtain a distribution of predictions for each population. **e) Classification performance for simulated evolutionary classes.** We obtain a high classification accuracy across evolutionary classes (86.1%). Positive and combination classes are predicted with 98.3% and 98.1%, respectively. We observe a reduction in accuracy in neutral (80.83%) and negative (81.25%) classes of evolution.

Here we use advanced statistical and deep learning techniques to study the underlying evolutionary mechanisms driving cellular dynamics in pre-cancerous and normal hematopoietic populations. We generated somatic variant calls from 92 individuals who subsequently progressed to AML (preleukemic cases), and 385 age- and sex-matched healthy controls from the European Prospective Investigation into Cancer and Nutrition (EPIC) study (7,17). Error corrected sequencing was performed on whole blood for 261 genes (xGen^®^ AML Cancer Panel) implicated in AML at approximately 5000X coverage and is described in detail elsewhere (7).

Our approach to estimate intra-evolutionary processes underscoring hematopoietic dynamics is outlined in ***Figure 1b-d***. Briefly, we consider sequencing data from each individual to be derivative of a hematopoietic stem cell population and extract population-level summary statistics to describe patterns of somatic mutations in the mature blood cell pool (***Figure 1b***). Summary statistics include counts of mutations, both overall and parsed according to whether they are silent or missense/nonsense, as well as summaries of variant allele frequency (18,19). Summary statistics, such as these and others, are frequently used to test for departures from neutrality in population genetics and, collectively, can be used to discriminate among mutation rates and selective pressures acting on polymorphisms segregating in genomic regions of interest (10,11,18–22). Using a deep learning classifier, trained on sequencing data simulated across range of realistic Wright-Fisher processes, we are able to classify individuals according to their dominant selection regime. We show how mutation rate and selection work in conjunction to influence the evolutionary trajectories of mutations in blood cell populations and highlight the critical role that mildly damaging mutations play in cancer prevention.

## Results

### Multi-task deep neural networks recover evolutionary dynamics and parameters with high accuracy

To infer evolutionary processes acting within each blood cell population, we trained an ensemble of deep neural networks (DNNs), hereafter “classifier”, using summary statistics derived from populations simulated across a range of evolutionary scenarios as input features, (***Figure 1b-d***). A total of 4.6 million simulations were produced and we used these to create a look-up grid of parameter combinations representing a comprehensive range of plausible evolutionary scenarios. Simulations were performed under a general Wright-Fisher model with four variable parameters: mutation rate, probability of a mutation being beneficial (*p*), coefficient of positive selection (*s_p_*; corresponding to the relative fitness advantage of cells with this mutation), and coefficient of negative selection (*s_n_*; relative fitness disadvantage of cells with this mutation) (23). The effects of selection were restricted to nonsynonymous sites and synonymous sites were considered to be neutral. Each unique combination of four parameters corresponds to a distinct evolutionary model, however, as an overall summary, each model can be collapsed into one of four overarching evolutionary classes as determined by the combination of selection coefficients (*s*): neutral (*s*= 0), positive selection only, negative selection only, and combination models which allow for the accumulation of mutations subject to both positive and negative selection within the same cellular population (see Online Methods). Each neural network within our ensemble classifies a population into one of the four evolutionary classes and estimates the four parameters comprising a given model (***Figure 1c***). Through comparing the outputs of our classifier, we can determine uncertainty in the classifications (***Figure 1d***) (24).

We were able to obtain a high classification accuracy for populations simulated under positive-only (0.98) and combination (0.98) models and relatively high accuracy for populations simulated under neutral (0.81) and negative-only (0.81) models when testing our classifier on a held-out test set of simulations for point mutations (10% of our simulated data) (***Figure 1e, Supplementary Figure 1***). A perturbation analysis of our inverse model showed that we can predict positive-only or combination evolutionary classes with high certainty but that our model has some difficulty distinguishing between neutral and negative-only selection when there were few mutations. Fewer mutations, or a lower level of variability in the population, could arise following a selective sweep or when populations are subject to a lower mutation rate (***Supplementary Figure 2***). Additionally, a lack of mutational information could be attributable to the selective removal of SNPs as a result of negative selection (***Supplementary Figure 3***). Distinguishing between neutral evolution and negative selection is a challenge itablen population genetics as weakly damaging mutations can segregate in the population at low frequencies and have a mild impact on reducing variability at linked loci (12,13).

### Hematopoietic populations show evidence of positive and negative selection regardless of disease outcome

We applied our classifier to preleukemic cases and healthy controls in order to infer population-level evolutionary processes. We find that in the majority of individuals (71%), both controls and cases, the hematopoietic population does not evolve neutrally (***Figure 2a***). We reject models of neutral evolution in the majority of cases (79%) and controls (64%) and we observe a significantly higher departure from neutrality in cases than controls (Pearson’s chi-squared test, *P* = 0.006). The majority of cases (62%) and the plurality of (43%) controls fit combination classes of evolution; in other words, we are able to detect signatures of both positive and negative selection in their blood biopsy indicating that the functional impact of passenger mutations cannot be overlooked. We observe that higher levels of predictive uncertainty correspond with a reduction in segregating mutations in the mature blood cell pool, in keeping with our classifier’s performance on simulated populations (***Supplementary Figure 4***). In summary, few cases or controls are evolving neutrally, and we find evidence of positive selection and of negative selection in the majority of preleukemic cases and healthy controls.

**Figure 2.**
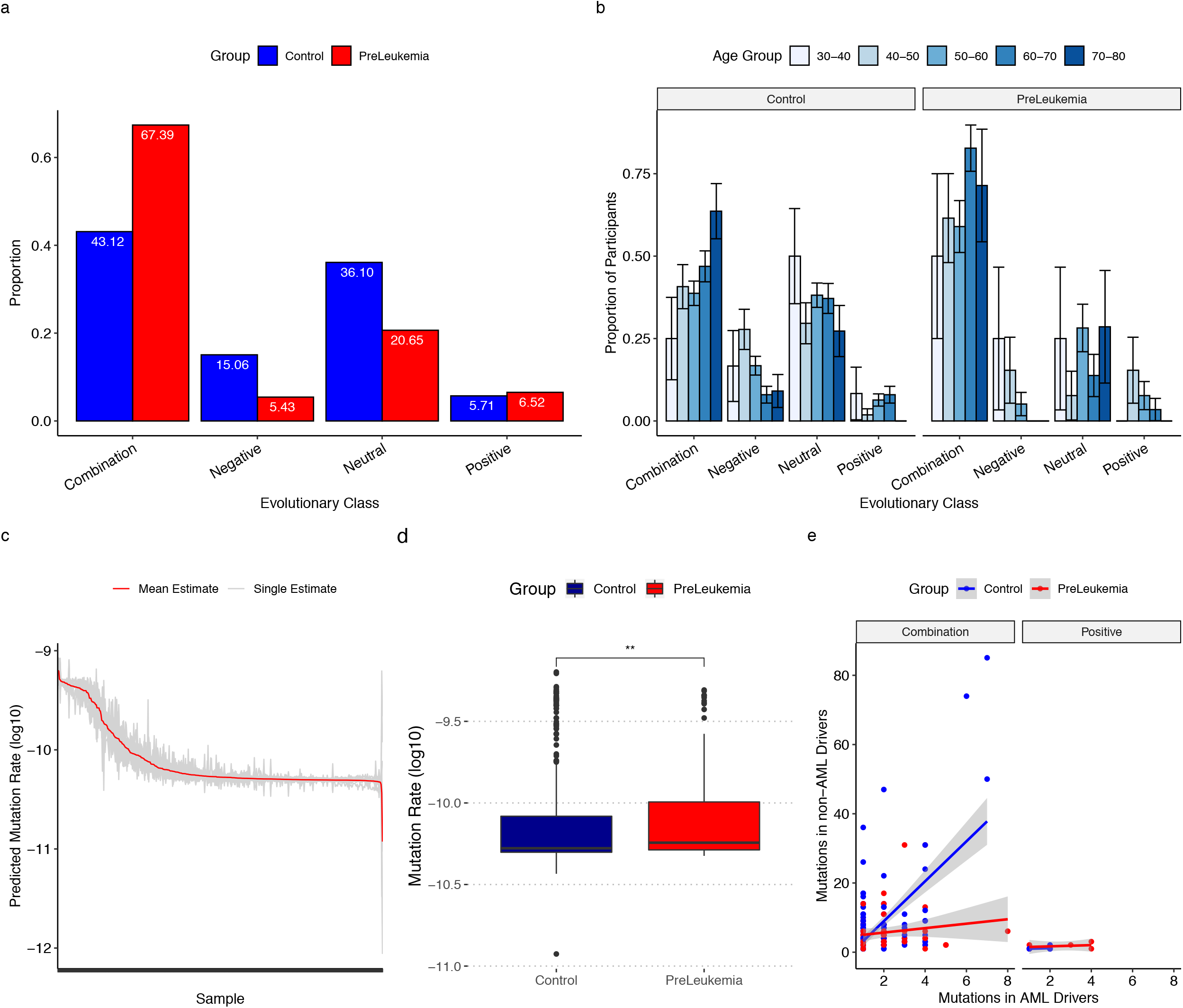
**a) Evolutionary classes in preleukemic and healthy blood populations.** The majority of blood populations do not evolve neutrally (72%). Similarly, only 9% of individuals fit positive models of evolution. Populations do not evolve neutrally in the majority of preleukemic cases (79%) and healthy controls (64%). The majority of preleukemic (62%) and the plurality of healthy (43%) individuals fit combination (both beneficial and damaging mutations arising) classes of evolution. **b) Age-associations across evolutionary class predictions.** Participants were stratified into 10-year age windows. We calculated the proportion of individuals within each age bin fitting each evolutionary class for preleukemic individuals and healthy controls. Standard errors for each proportion were calculated by p(1-p)/n where p is the proportion of individuals fitting a particular class and n is the total population. **c) Range of mutation rate estimations across cohort of participants.** We show the estimated mutation rate for each sample from each DNN in our classifier (grey). The mean estimate from the classifier outputs is shown in red. Mutation rates (y-axis) are log-transformed and scaled to a population size of 10,000. **d) Preleukemic blood populations have a higher mutation rate than healthy controls.** The best fit mutation rate is plotted for each individual grouped according to outcome status (Control = blue, preleukemic=red). Preleukemic cases are found to have a modest yet significantly higher mutation rate than controls (Wilcoxon rank sum test, *P*= 0.0045). **e) Relative passenger to driver mutation proportion across evolutionary classes.** The number of mutations in known driver genes is plotted against the number of mutations in non-driver genes for each individual blood population with healthy controls shown in blue and preleukemic individuals in red. A linear regression was performed to approximate the ratio of mutations within each outcome group. In combination models, controls have a significantly higher proportion of mutations in non-AML driver genes compared to drivers compared to preleukemic cases (Control: β=5.765; Case: β=0.642; F test, p <0.0001) We are not able to capture this same association in the positive-only class (Control: β=0.167; Case: β= 0.174; F test, p = 0.98).

### Hematopoietic populations evolve in an age-dependent manner

As ARCH is known to be an age-associated phenomenon, we investigated if there is any association between the age of an individual and the selective pressures governing hematopoietic dynamics. Participants were binned into age groups spanning ten-year intervals and the proportion of participants in each agerange fitting each evolutionary class was calculated. We find clear associations with age and the dominant class of evolution in individuals (***Figure 2b***). Specifically, we observe an age-related decline in the proportion of controls fitting negative-only or neutral classes of evolution, and a parallel increase in controls fitting the combination class. Our results are consistent with a model in which individuals accumulate passenger mutations as they age, some of which will have a slightly damaging effect. In parallel, with increasing mutation accumulation, there is an increased likelihood of a rare driver event occurring which would cause an individual to shift to a combination, or positive-only, class of evolution. We find that many preleukemic cases show evidence of positive selection at a younger age than controls. In particular, in preleukemic cases, where age-associated clonal expansions have been previously reported, we observe an increased overall proportion of individuals fitting combination models in younger age groups indicating that driver events have occurred earlier. At a young age, driver mutations are likely to be arising on a background with fewer passenger mutations and thus may experience a relatively higher fitness advantage compared to the same mutation arising on a background with a greater number of passenger mutations; a hypothesis that we investigate in the following section.

### Controls have a higher proportion of passenger-to-driver mutations than cases

To investigate if the proportion of mutations in known driver genes compared to non-driver genes could explain some variation in outcomes, we compared the types and patterns of mutations between the cases and controls. We annotated mutations as drivers if they occurred in driver genes found to be highly mutated in the Cancer Genome Atlas Acute Myeloid Leukemia project (***Supplementary Figure 5***). We first asked whether cases simply had more mutations, thus predisposing their blood populations to cancer. Consistent with previous reports (7), our cases had more mutations on average than age-matched healthy controls and, in the positive-only class of evolution, more mutations in known driver genes (***Supplementary Figure 5***) (Wilcoxon rank sum test, p value = 0.0027). A higher mutation count in cases is in keeping with our classifier’s prediction that there is a modestly higher mutation rate in a preleukemic context (mean mutation rate: μ = 1.191368e-10 per bp per division) compared to healthy controls (mean mutation rate: μ = 1.094472e-10 per bp per division) (Wilcoxon rank sum test, p value = 0.0045) (***Figure 2c-d***). We estimated mutation rate assuming a population size of 10,000 and we have scaled our estimates to account for varying estimates of HSC population size (***Supplementary Figures 6-8***).

Intriguingly, we do observe mutations in driver genes in healthy controls in the presence of positive selection. It is possible that these individuals do not progress to disease if the mutations do not alter clonal growth dynamics or driver gene function, or if driver mutations are arising in competing clones. However, another possible explanation for the differences in outcome is the proportion of driver to passenger mutations. We find that in controls, there is a higher proportion of passenger-to-driver mutation ratio in combination models (F-test, *P* <0.0001) (***Figure 2e***). We do not see this difference between cases and controls in the positive-only modes of evolution (F-test, *P*=0.98), but our sample size in this comparison is low, so our power to detect such a difference may be insufficient. The increased proportion of mutations in passenger genes compared to driver genes in the combination class is consistent with a model in which negative selection acting on passenger mutations is playing a protective role in inhibiting or stalling clonal expansions.

### Distinct patterns of inferred pathogenicity associate with evolutionary classes

To determine whether some passengers are playing a protective role, we scored each mutation according to how likely it was to affect protein function and conservation after blood sample classification. In doing so, we are able to independently evaluate the performance of our evolutionary predictions. We scored mutations using the Combined Annotation-Dependent Depletion (CADD) algorithm (***Figure 3a***) (25). Using a combination of functional prediction, conservation, epigenetic measurements, gene annotations and the sequence surrounding a given variant, CADD provides a measure of the functional impact of single nucleotide variants, and small insertions/ deletions, in the genome. CADD scores assess whether a mutation alters protein function, and have difficulty distinguishing between whether it changes protein expression, inhibits its activity, or causes the protein to be constitutively active; as such we will call mutations with high CADDs “function-altering”.

**Figure 3.**
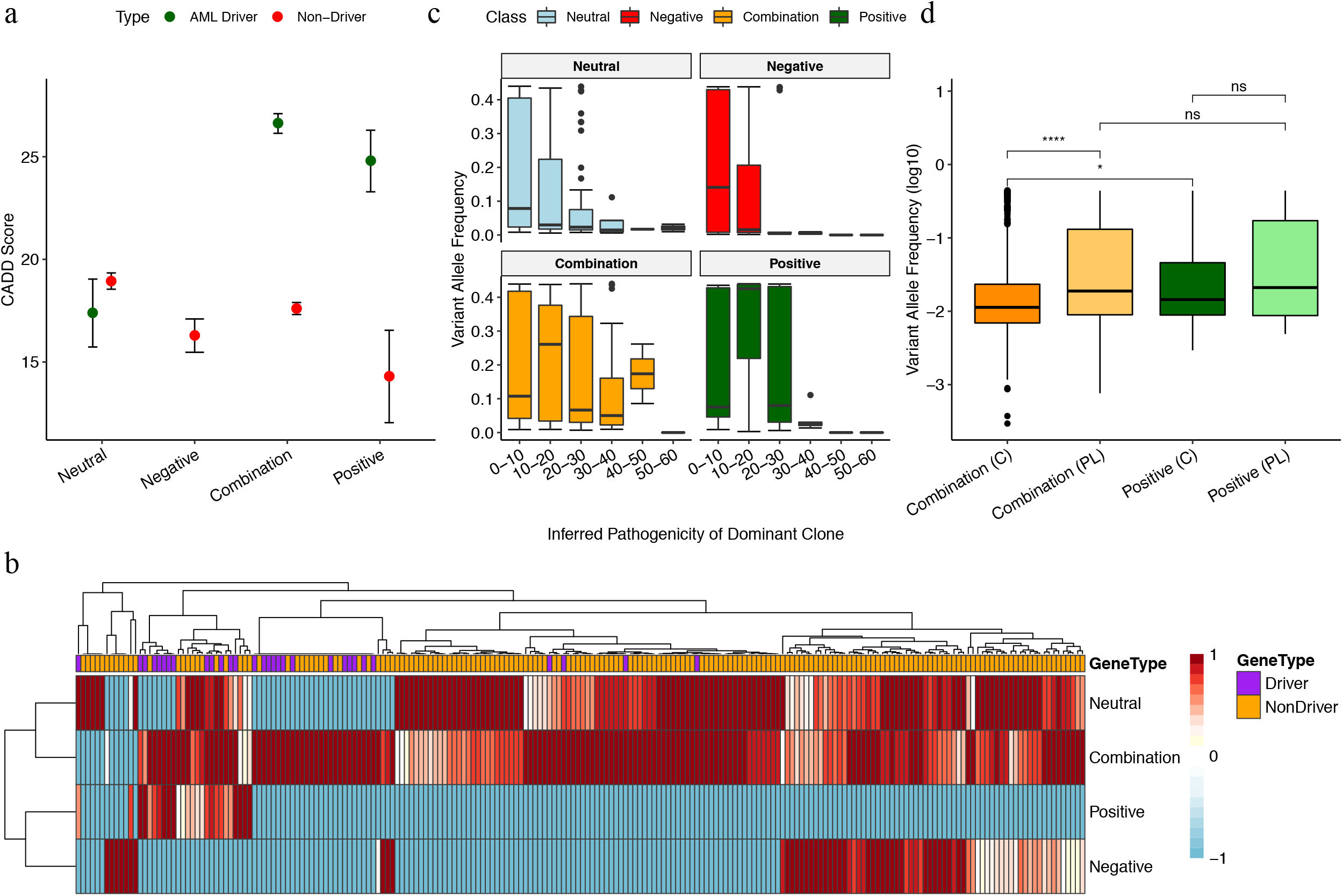
**a) Predicted functionality of mutations in each evolutionary class.** Average CADD scores were calculated for mutations in known driver genes and non-driver genes. We capture a significant (Wilcoxon rank sum test, beneficial models: *P* = 0.0016; combination models: *P* <2e-16) enrichment of high CADD scores in driver genes compared to non-driver genes. We do not observe a significant difference between CADD scores across mutations in driver genes and non-driver genes in neutral classes (*P*= 0.3). The average CADD score assigned to passenger mutations in negative models is significantly lower (Wilcoxon rank sum test, *P* = 0.004) than passenger mutation CADD scores in the neutral class. **b) Genes enriched for functional mutations across evolutionary classes.** The mean CADD score of mutations in each gene was calculated for each evolutionary model. Average scores were normalized across models and genes not harbouring mutations were set to −1 (blue) to differentiate between a lack of information and CADD scores of 0 (white). Values closer to 1 (dark red) indicate that genes (columns) are highly enriched for functional mutations within a particular evolutionary class (rows). A panel above the heatmap indicates if genes are known AML driver genes (purple) or non-driver genes (orange). **c) Inferred pathogenicity of dominant clone is correlated with the variant allele frequency across different evolutionary classes.** We isolated the dominant clone within each individual blood pool. The CADD scores of each clone was binned into intervals of 10 and the distribution of variant allele frequencies was calculated for each bin. We observe a wider distribution of CADD scores in neutral and combination models. In positive classes of evolution, clones are found at a higher frequency suggesting that they sweep to fixation. Similarly, in negative models of evolution, we observe a surplus of clones in the low CADD score bins which appear to segregate at a lower frequency. **d) Impact of Negative Selection on Clonal Expansions.** We investigated if negative selection acting on passenger mutations impacted clonal expansions in a healthy and preleukemic context. To do so, we plotted the log transformed variant allele frequency (VAF) of mutations found in cases and controls predicted to be evolving in a positive class (green) and combination (orange) classes. VAF distributions were plotted separately for preleukemic individuals (light green/ light orange) and healthy controls (dark green/ dark orange). We find that we are not able to discriminate between VAF distributions of mutations in healthy and preleukemic individuals in the positive class. Further, we are not able to discriminate between positive models of evolution and preleukemic individuals who fit combination models of evolution. However, we find that controls fitting combination classes of evolution have a significantly lower VAF distribution compared to both preleukemic cases fitting combination models (Wilcoxon rank sum test, *P* < 2e-16) and controls fitting positive models (Wilcoxon rank sum test, *P* = 0.02).

Overall, we observe that mutations falling in known driver genes tend to have a higher CADD score than mutations in non-driver genes. However, in keeping with our expectations of neutral evolution, we do not observe a significant difference in CADD score between mutations in known driver genes and non-driver genes in neutral cases (Wilcoxon rank sum test, *P* =0.3), suggesting that these mutations are not function-altering and confer no relative fitness advantage or disadvantage to the clone in which they are found. In comparison, in individuals showing evidence of positive selection (positive and combination), mutations in known driver genes had significantly (Wilcoxon rank sum test, positive models: *P* = 0.0016; combination models: *P* < 2.2e-16) higher CADD scores than mutations in non-driver genes. Further, we observe that the average CADD score assigned to passenger mutations in negative-only models is significantly lower (Wilcoxon rank sum test, *P*= 0.004) than passenger mutation CADD scores in the neutral class suggesting that negative selection plays a role in removing the more damaging mutations and decreasing the overall pathogenicity of segregating mutations. The role of negative selection in decreasing the overall pathogenicity of the blood pool is further supported by the average CADD scores of passenger mutations in the combination class being greater and smaller than the average score of passenger mutations in the negative-only and neutral class, respectively. In the absence of recombination, mutations which would typically be removed are able to continue to segregate in the blood population in the presence of positively selected driver mutations and, accordingly, we observe a higher average pathogenicity of passenger mutations in the combination class. Finally, it is worth noting that the passenger mutations in the positive-only class have a significantly lower pathogenicity than those in the combination class. Passenger mutations with a higher pathogenicity are likely to be subject to stronger negative selection thus conferring a protective effect to the individual in the presence of positive selection acting on drivers. A better understanding of how these potentially protective mutations are distributed across genes would allow us to identify which genes might be critical in preventing clonal expansions.

### Clusters of genes are enriched for functional mutations across evolutionary classes

To investigate if certain genes are more frequently found to play a protective role when mutated, we used a hierarchical clustering approach to ascertain which genes are enriched for function-altering mutations in each evolutionary class (***Figure 3b***). Each gene was associated with a score vector whose elements were the mean CADD score per mutation in evolutionary class. We find that there are distinct subsets of genes which are enriched for function-altering mutations in the presence of positive and negative selection alone. Reassuringly, we find that many known driver genes (*DNMT3A, TET2, IDH2, TP53*) are enriched for functional mutations among positive and combination classes of evolution only and not among neutral or negative classes of evolution (***Supplementary Figure 9***). Further, we observe that there is an overlap of genes enriched for functional mutations in negative-only and combination classes of evolution, and not in neutral or positive-only models of evolution indicating that these genes might experience stronger negative selection.

We next asked if the inferred pathogenicity of mutations in the dominant clone corresponds with the frequency at which it is observed in the mature blood cell pool. To do so, we evaluated the relationship between CADD score and the frequency of the dominant clones, defined as the clone with the highest variant allele frequency in an individual, in each class (***Figure 3c***). We find that clones are able to rise to fixation in the absence of both negative and positive selection where the primary driving force of evolution is genetic drift. Clones rising to a high frequency stochastically could, in part, be explained by a reduction in the effective population size of the HSC population owing to a small population of stem cells with a higher fitness dominating blood cell production. With a reduced population size, mutations are able to rise to higher frequency and become fixed in a population more rapidly. However, only mutations with a low CADD score are found at high frequencies in the neutral class. As expected, in the presence of negative selection, we observe a depletion of clones in the higher pathogenicity categories as they have likely been removed by selection. Clones which persist in the negative-only model could indicate a functional threshold at which mutations are tolerated and not removed by selection. Conversely, in the positive-only class, clones, including those with a high pathogenicity, are found at higher frequencies. We observe a higher variance in CADD scores in the combination class which is consistent with our expectation that, when neither positive or negative selection are able to act efficiently, variants will segregate at intermediate frequencies rather than sweeping to high fixation or being purged from the population, respectively.

### Selective interference may be associated with slowing clonal expansions

Having established that in the combination class, controls have significantly higher non-driver to driver ratios, and that these non-drivers have significantly higher CADD scores than those in the positive-only class, we then ask whether non-drivers played a role in preventing progression to AML through selective interference. Selective interference is particularly relevant as studies report that driver mutations, while found in both healthy controls and preleukemic cases, tend to segregate at a much higher frequency in a preleukemic context (7). We propose that selective interference, where linkage between sites under multiple selective pressures will define the overall impact of selection acting on the population, could play a role in preventing mutations from rising to a high frequency in controls either through passenger mutations hitchhiking within the same clones as driver mutations, or if driver mutations arise in different clones and are competing for dominance in a finite cell pool (13,26,27). We expect that clones under purely positive selection will be found at higher frequencies in blood compared to those subject to a combination of positive and negative selection where interference might play a role in preventing selective sweeps.

We find that mutations in preleukemic cases fitting a combination class tend to segregate at a significantly higher frequency compared to controls (Wilcoxon rank sum test, *P* < 2.2e-16) (***Figure 3d***). However, we do not observe a difference in the frequency at which mutations are found to segregate between cases and controls fitting positive models of evolution or between cases fitting combination and positive models of evolution. Decreased variant allele frequencies in healthy individuals fitting combination models is consistent with our prediction that selection acting on a subset of mutations in healthy controls prevents progression to disease even in the presence of positive selection. However, we do observe signatures of negative selection in preleukemic contexts which suggests that the impact of selection acting on passenger mutations, while detectable through our methods which incorporate multiple summaries of the data, remains negligible with respect to clonal progression. Further, we do observe that, while not significant owing to sample size, preleukemic cases fitting combination models tend to have a later age of diagnosis than preleukemics fitting positive-only evolutionary classes indicating that negative selection might play a role in slowing progression to disease. Our findings suggest that not all passenger mutations are equal in that some might be more efficient in preventing disease-associated clonal expansions. Further, through accounting for mutations which segregate alongside driver mutations, we would be able to greatly improve our understanding of ARCH as a biomarker for disease and better predict who is at risk of progressing to cancer.

## Discussion

Our ability to determine the evolutionary processes governing the impact of age-associated mutations on hematopoietic fitness is dependent on the deep interrogation of blood profiles and our ability to discriminate amongst alternative evolutionary models. Estimates of cellular evolutionary features are dependent on simultaneous inference of mutation and selection. Most tests to detect selection are based on the null-hypothesis that mutant alleles are selectively neutral and have proven to be highly successful in detecting positive selection in germline and somatic contexts. However, the detection of negative selection in populations has remained challenging as they are reliant on the capture of rare variants which, prior to deep coverage sequencing, are typically not reliably detected in cellular populations (12). Many tests of neutrality, including *dN/dS* ratios which are often used to study selection in paired tumour normal samples, are only sensitive to a conservative range of selection parameters and can be confounded by the timing of events. However, determining the mode of selection acting on a mutation is a key parameter to understanding tumour evolution as it offers critical insight into the evolutionary fate of the mutation in the population. In a somatic context, this becomes increasingly more important as negative selection acting on mutations co-occurring in the same clone as positively selected driver mutations could significantly alter the evolutionary trajectory of the mutant expected under positive or neutral evolution alone.

Using newly-developed population-genetic neural-network approaches which exploit the impact of linkage in clonal systems and combinations of informative summaries of mutation data (12,13), we are able to discriminate signatures of negative selection from neutrality. Our methods are able to classify individual liquid biopsies as evolving under different selective regimes and enable us to evaluate the combined impact of both positive and negative mutations on patterns of clonal dominance in the mature blood cell pool of healthy and precancerous individuals. We find that hematopoietic populations largely do not evolve neutrally and that the presence of negative selection acting on mutations in non-driver genes plays an important role in disease development across aging blood systems. In line with other studies, we find that we do observe ARCH occurring in blood populations presumed to be evolving strictly under neutral evolution, however in most instances, the trajectory of clonal mutations in the blood appears to be governed by the complex interplay of both positive and negative selection (28). Further, we provide some of the first estimates of the rate at which mutations accumulate the aging hematopoietic system in both healthy and premalignant contexts. The rate at which ARCH associated mutations occur in healthy and premalignant contexts has, to our knowledge, not been previously characterized owing to genomic and statistical challenges.

Our approaches untangle interacting and sometimes confounding factors in somatic evolution. We captured an important phenomenon within blood: that damaging mutations accrue along with ARCH-associated driver mutations. Indeed, these damaging mutations may hinder the rate that clones drive through the population. The presence of negative selection acting on passenger mutations in the presence of driver mutations offers a potential explanation for why some individuals who harbour driver mutations do not progress to disease. However, negative selection appears to play an important role in lowing the overall pathogenicity of the mature blood cell pool, thus maximizing the fitness of the individual; a finding which suggests that there could be an advantage to retaining mildly damaging mutations in cellular populations if they confer a protective effect in in the presence of a driver mutation. Future work with long-read sequencing and single-cell sequencing is required to experimentally phase somatic mutations genome-wide will help determine the timing of such mutations and whether such mutations are occurring in the same or different clones and the implications on mutational trajectories.

## Online Methods

### Sequencing and data processing in EPIC Cohort

Participant selection, sequencing and mutation calling have been described in detail elsewhere (7).

### Calculation of Summary Statistics

ummary statistics were calculated per individual with sequencing data from each individual considered to be derivative of an HSC population. We calculated a total of 16 summary statistics for each hematopoietic population including population genetics statistics used to describe the site frequency spectrum (Tajima’s D (18) and Fay and Wu’s H (19)), counts and ratios of synonymous and nonsynonymous mutations both overall and for low (>0.1), intermediate (0.1-0.8) and high (>0.8) variant allele frequency windows, and the number of mutations in known driver genes.

### Simulations of hematopoietic populations across evolutionary scenarios

Hematopoietic populations were simulated as clonal haploid populations evolving as a Wright-Fisher process forward in time using the software SFScode (23). We simulated haploid populations with an effective population size of 10,000 and a null recombination rate. A 1Mb region of the genome with 1:2 ratio of synonymous and non-synonymous positions were simulated, where synonymous sites are not subject to either negative or positive selection. We simulated a grid of evolutionary scenarios across a plausible range of parameters. Selection coefficients affecting each nonsynonymous mutation are sampled from a gamma distribution (29). Owing to variation associated with cancer evolutionary parameters, we simulated ranges of mutation rates (from 1e-7 to 1e-5 mutations per base pair per generation), proportions of beneficial mutations (from 0 to 1; 0 being that all mutations are deleterious and 1 being that all mutations are beneficial), and rate parameters for the gamma distribution from which selection coefficients are sampled (from 0.001 to 0.005). The shape (*α*) and rate (*β*) parameters of the gamma distribution were also varied. Effectively, in a subset of simulations, the rate parameter for the gamma distribution of negative selection coefficients range from 0.001 to 0.02475. We compute the means of selection coefficients based on the rates and shapes of the tested gamma distribution (μ= *α*/*β*). For each set of parameters, we perform 2000 replicates. Additionally, we simulated populations evolving under neutral models of evolution (no selection). Summary statistics for simulated populations were calculated as described for the EPIC participants. The number of mutations subject to positive selection in each simulation is a function of the probability of a mutation being beneficial (*p*) and the number of nonsynonymous mutations. Each simulation was classified into one of four overarching evolutionary classes if the following conditions were met:

| **Class** | **$\mu$** | **$p$** | **$s_p$** | **$s_n$** |
| --- | --- | --- | --- | --- |
| **Neutral** | All | 0 | 0 | 0 |
| **Positive Selection** | All | $0 < 0.5; 1$ | $0 < x < 0.05$ | 0 |
| **Negative Selection** | All | 0 | 0 | $0 < x < 0.05$ |
| **Combination** | All | $0 < x < 1$ | $0 < x < 0.05$ | $0 < x < 0.05$ |

### DNN Ensemble

We collated summary statistics from each simulation to create our main training dataset of populations evolving under different selection pressures. Our simulations were subdivided into training (90%) and test (10%) sets. To create a unique training set for each instance of our DNN ensemble, we further subsampled our training set into 10 sets consisting of 250,000 unique evolutionary models. Sampling was done with replacement and the number of models in each class was downsampled to match the minority class to avoid class imbalance. We trained a total of ten deep neural networks (DNNs) independently. Each DNN was trained as a multi-task neural network in order to both classify a population into one of four overarching evolutionary classes (positive, negative, combination or neutral) and predict four continuous parameters (mutation rate (μ), the probability of a mutation being beneficial (*p*), coefficient of positive selection, and coefficient of negative selection). A benefit of our ensemble-based approach is that, for each blood cell population, each DNN emits a softmax probability distribution across the four overarching evolutionary classes. In a conventional classification task, the class with the highest probability will be selected as the best fit. However, as we are employing an ensemble-based approach, we are able to obtain a distribution of predictions for each population so as to measure the uncertainty associated with each prediction. For each individual, we calculated the mean and standard error for each softmax probability across the four evolutionary classes and accepted the class with the maximum softmax probability as the class of best fit. Similarly, for each regression task we calculate the mean and standard error for each parameter across the predictions for each individual.

### Predicting best fit class and estimating parameters for EPIC individuals

To obtain the best fit evolutionary class for each individual, we calculated the mean and standard error for each softmax probability across the four evolutionary classes and accepted the class with the maximum softmax probability as the class of best fit. To calculate age distributions of evolutionary classes, we bin participants into 10-year age windows. We calculate the proportion of individuals within each age bin fitting a particular evolutionary class for preleukemic individuals and healthy controls. Standard errors of each proportion were calculated by 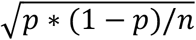 where *p* is the proportion of individuals fitting a particular class and *n* is the total population. We calculate the mean and standard error for parameter predictions to estimates for mutation rate and the probability of a mutation being beneficial.

### Predicted fitness effects of mutation

Scaled CADD scores for mutations in each evolutionary model were obtained using the online variant annotation tool (25). Average CADD scores were calculated for mutations in known driver genes and known passenger genes and used to independently validate mutational dynamics in each model. To identify genes enriched for mutations in different functional classes, the mean CADD score of mutations in each gene was calculated for each overarching evolutionary model. Average gene scores were normalized across models through taking 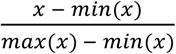 and genes not harbouring mutations were set to −1 to differentiate between a lack of information and CADD scores of 0.

## Acknowledgements

We thank the Ontario Institute for Cancer Research Scientific Advisory Council (Christina Curtis and Elaine Mardis) for thoughtful comments on the study. Support was provided by the Ontario Ministry of Research and Innovation to PA and KS, a CIHR Frederick Banting and Charles Best Canada Graduate Scholarship-Masters and Terrence Donnelly Center Cecil Yip Award to KS, and the Vector Institute for Artificial Intelligence to KS and QM. Genomic data generation and analyses was supported by the Canadian Data Integration Center (CDIC) with funds provided by the Government of Ontario and the Government of Canada through Genome Canada and Ontario Genomics (OGI-136).

## Author Contributions

PA conceived of the study and PA, SW, QM and KS designed the analyses. LS, JD, and SA performed experiments and contributed data. AMH performed all simulations. KS, DS, AMH, BL, VB, QM, and PA performed all analyses. KS, QM, and PA wrote the manuscript.

