## Supplementary Figures for "Interacting evolutionary pressures drive mutation dynamics and health outcomes in aging blood"

**Supplementary Material**

**Supplementary Figure 1.** **Uncertainty associated with class predictions in EPIC cohort.** A benefit of our ensemble-based approach is that, for each blood cell population, each DNN emits a softmax probability distribution across the four overarching evolutionary classes. In a conventional classification task, the class with the highest probability will be selected as the best fit. However, as we are employing an ensemble-based approach, we obtain a distribution of predictions for each population so as to measure the uncertainty associated with each prediction. To obtain the best fit evolutionary class for each individual, we calculated the mean and standard error for each softmax probability across the four evolutionary classes and accepted the class with the maximum softmax probability as the class of best fit. **A)** Here, we show the mean (dark blue) and standard error (light blue) softmax probability for all EPIC participants. We find that for approximately half of the participants (n= 215 (51.5%)) we obtain average probability distributions of over 99%, with a maximum standard error of 0.008, indicating that we are able to predict overarching evolutionary classes with high certainty. **B)** All participants predicted with a high degree of certainty are classified as evolving under either beneficial or mixture models of evolution. Participants classified as evolving under negative or neutral classes of evolution typically exhibit increased levels of predictive uncertainty.

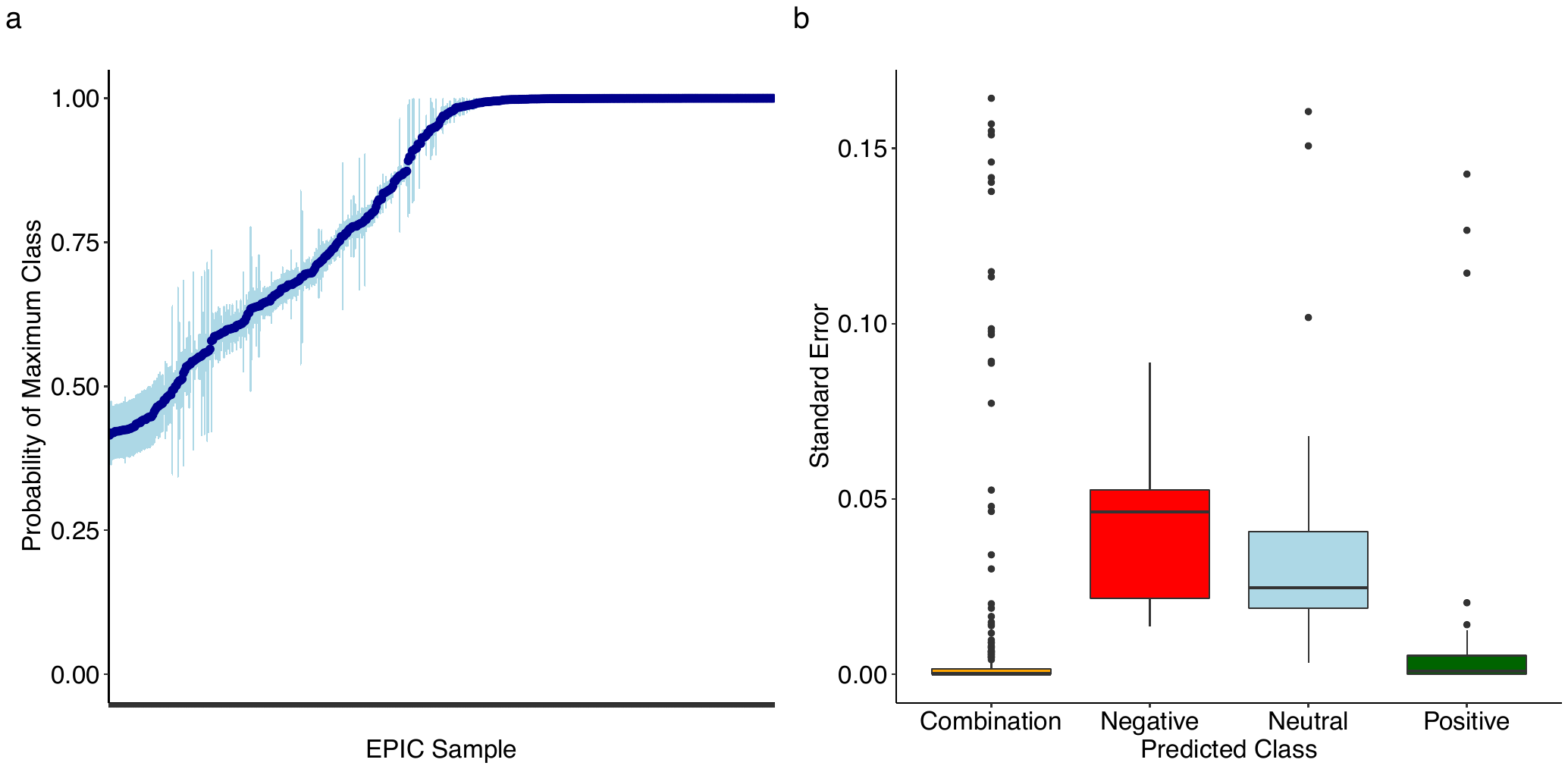

**Supplementary Figure 2. Impact of parameters on predictive accuracy.** We evaluated the distribution of parameters under which simulations were performed for neutral and negative classes of evolution. We show the proportion of the parameters where the evolutionary classes were correctly classified (green) compared to the proportion of parameters where incorrectly classified (red). Note, we only considered mutation rate and the co-efficient of negative selection as the other two parameters are not present in the neutral and negative classes (probability of a mutation being beneficial, and coefficient of positive selection). We observe an increase in the proportion of correctly classified populations in simulations performed with a higher mutation rate (left panel). Similarly, simulations performed with a lower mutation rate are more commonly misclassified. This is in keeping with our expectation that populations simulated with a lower mutation rate will have fewer mutations which corresponds with a decrease in information to inform the DNN ensemble. We do not observe an obvious trend in the proportion of simulations misclassified across co-efficients of negative selection.

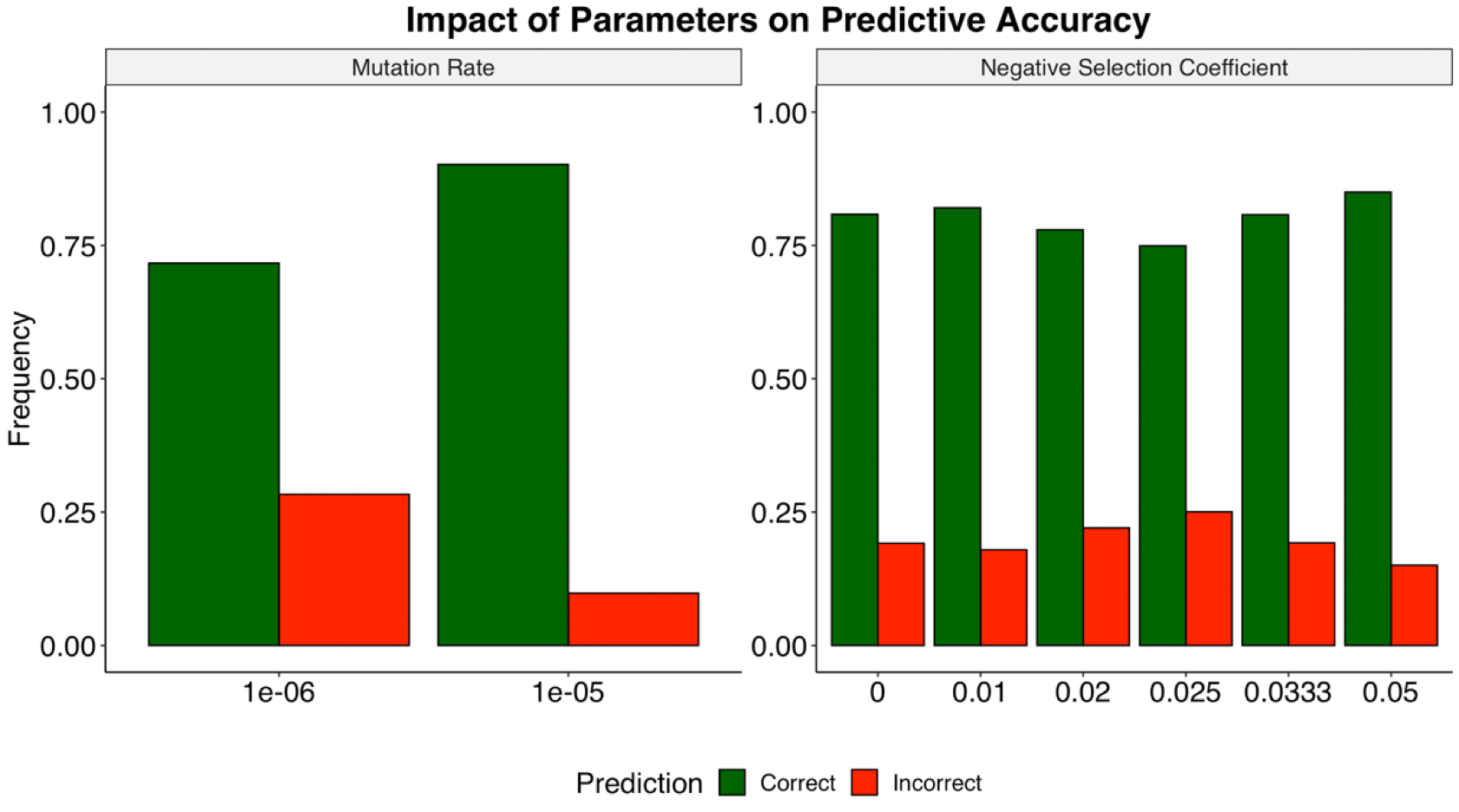

**Supplementary Figure 3. Impact of cumulative selective effect on accuracy.** We investigated the effects of the coefficient of negative selection on our ability to accurately classify neutral and negative classes of evolution. Specifically, we explored the cumulative selective effect, the number of mutations subject to a given selection coefficient, in our simulated populations. Cumulative selective effect was calculated by taking the product of the number of nonsynonymous mutations in a population and the coefficient of negative selection for each simulated population. We find that in the instances where we are able to correctly classify our simulated populations (green), we observe a higher cumulative selective effect compared to the instances where we are not able to correctly classify our simulated populations (red). This is in keeping with the expectation that the number of mutations segregating in a population is critical to informing evolutionary class predictions.

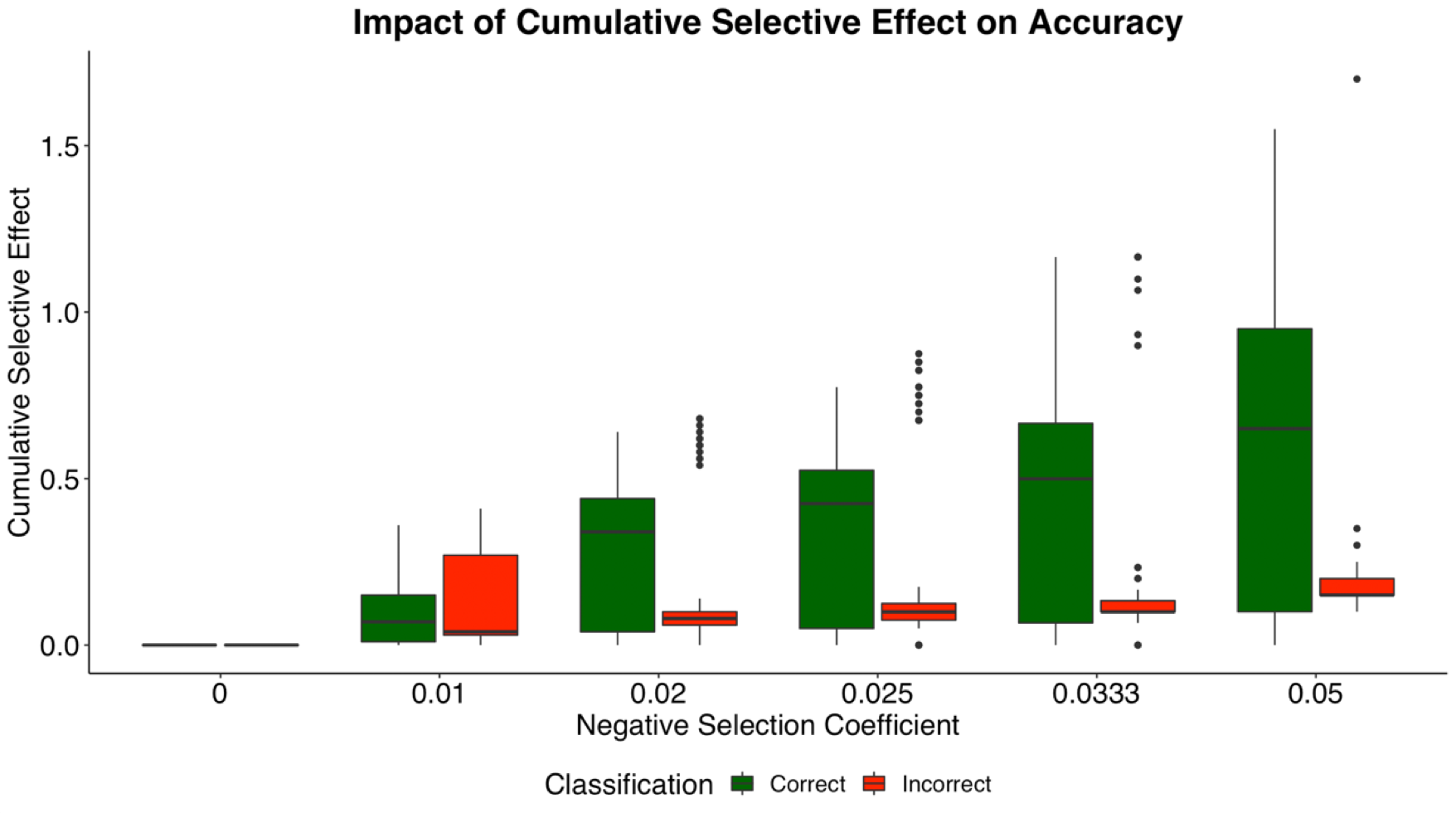

**Supplementary Figure 3. Impact of cumulative selective effect on accuracy.** We investigated the effects of the coefficient of negative selection on our ability to accurately classify neutral and negative classes of evolution. Specifically, we explored the cumulative selective effect, the number of mutations subject to a given selection coefficient, in our simulated populations. Cumulative selective effect was calculated by taking the product of the number of nonsynonymous mutations in a population and the coefficient of negative selection for each simulated population. We find that in the instances where we are able to correctly classify our simulated populations (green), we observe a higher cumulative selective effect compared to the instances where we are not able to correctly classify our simulated populations (red). This is in keeping with the expectation that the number of mutations segregating in a population is critical to informing evolutionary class predictions.

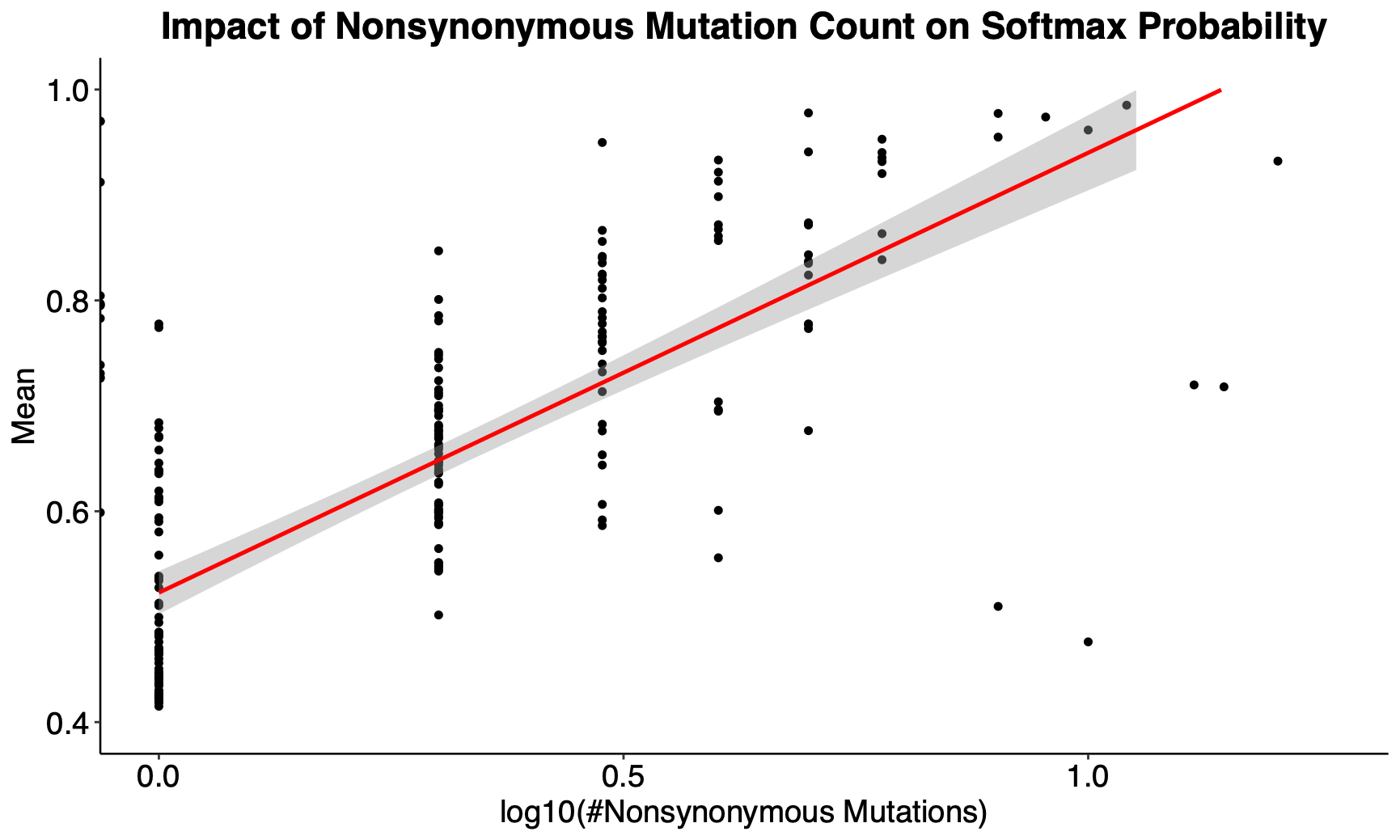

**Supplementary Figure 5. Mutational burden across evolutionary classes.** We investigated if there are higher numbers of passenger mutations in healthy individuals compared to controls. Genes were annotated as driver or non-driver genes based on if they were listed as one of the 32 driver genes in The Cancer Genome Atlas Acute Myeloid Leukemia project (DNMT3A, NPM1, FLT3, TET2, RUNX1, IDH2, TP53, IDH1, CEBPA, NRAS, WT1, KIT, PTPN11, KRAS, U2AF1, STAG2, PHF6, ASXL1, RAD21, EZH2, KDM6A, DIS3, SUZ12, CUL1, BCOR, NF1, THRAP3, CHD4, PRPF8, EGFR, MED12, CBFB). A) The mean driver and non-driver mutation count was calculated across healthy and preleukemic individuals fitting each evolutionary class. We observe significantly higher driver mutation counts in preleukemic individuals fitting positive and combination models. Further, we observe no driver mutations in individuals fitting negative models, preleukemic cases fitting neutral evolution, and only a slight increase in the average number of driver mutations in neutral controls. In almost all classes of evolution, with the exception of positive, we observe a higher number of mutations in non-driver genes in controls compared to preleukemic individuals.

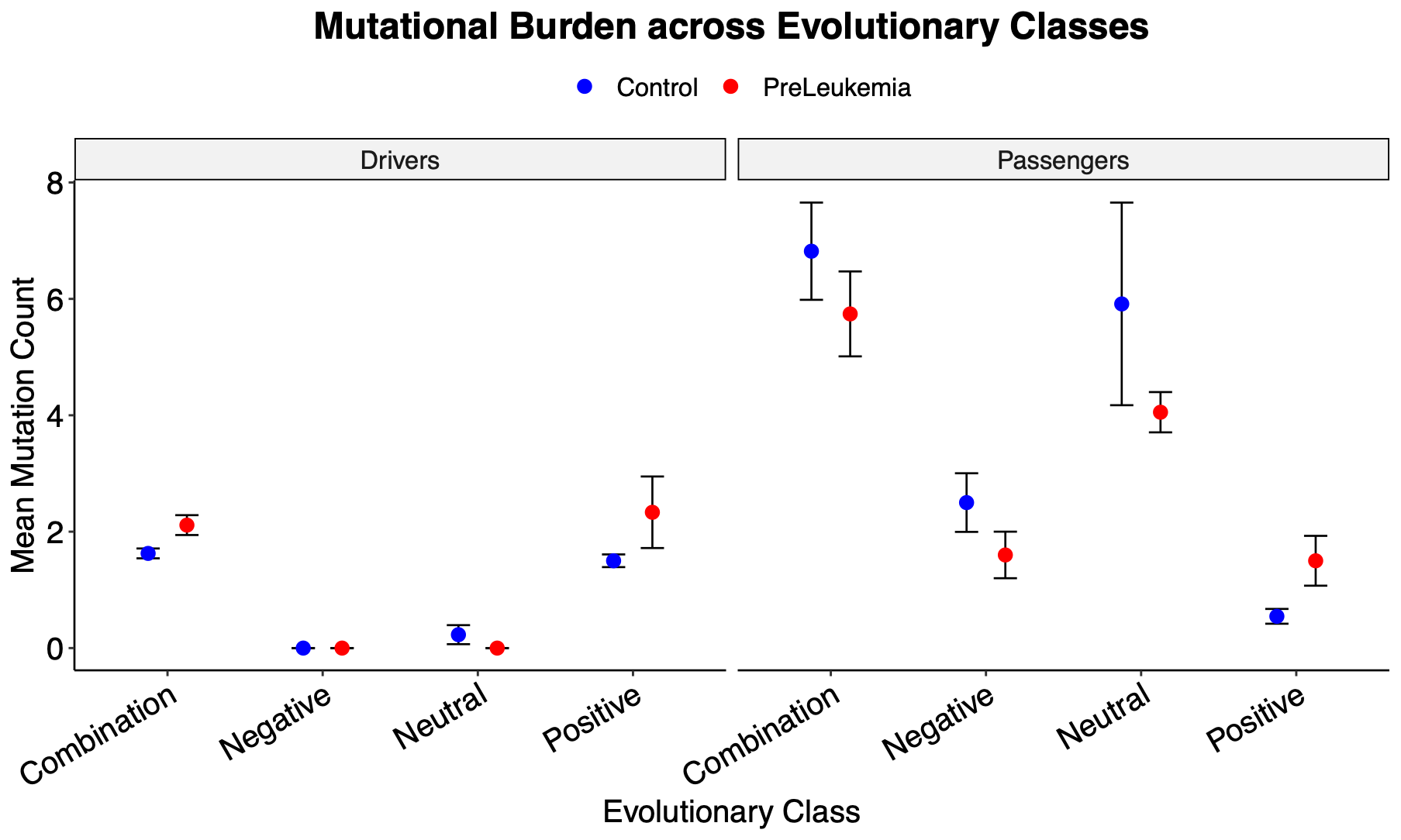

**Supplementary Figure 6. Distribution of mutation rate predictions for simulated data.** Density distribution of the mutation rate predictions for each true simulated mutation rates (dashed red line). Mutation rates are scaled to a population size (N) of 10,000. Mutation rate is plotted on a log10 scale and scaled to the per generation mutation rate. The means of the distribution of estimates for each mutation rate were relatively close to the true simulated mutation rates with the true mutation rate falling within one standard deviation of the mean estimate in both cases (true mutation rate = 4.98e-11 per bp per division, mean estimated mutation rate = 6.06e-11 ± 3.86e-11; true mutation rate = 5.01e-10 per bp per division, mean estimated mutation rate = 4.59e-10 ± 1.00e-10).

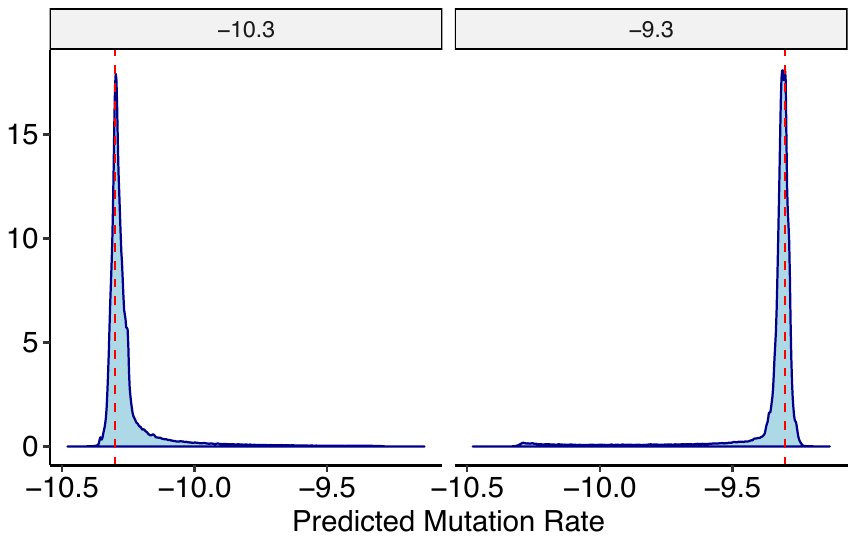

**Supplementary Figure 7. Mutation rate predictions across ensemble of DNNs.** Mutation rate prediction for each sample (x-axis) is shown for each DNN within the ensemble (y axis)**.** Mutation rates vary by two orders of magnitude across the cohort regardless of health outcome.

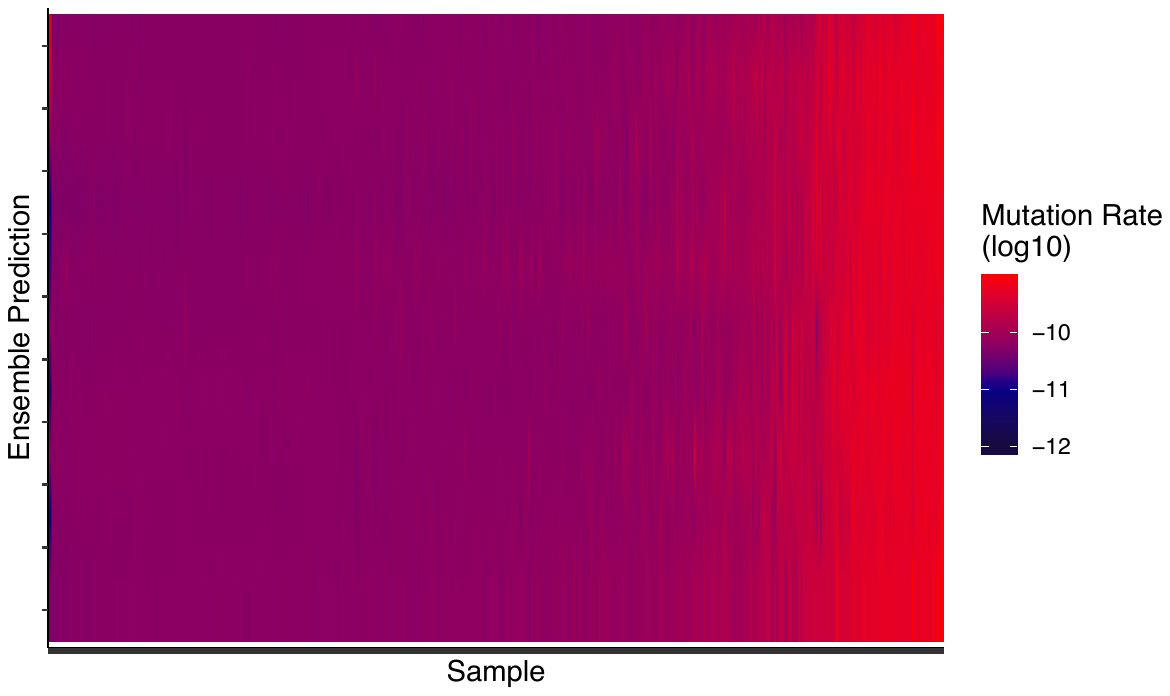

**Supplementary Figure 8. Impact of population size on mutation rate.** The expected number of mutations in a population (𝜃) is a product of the population size *N* and the per generation mutation rate (μ): 𝜃=4*N*μ. Estimates of mutation rate can be extended to account for the range of population size estimates (10,000 – 200,000) which exist for the hematopoietic stem cell population. For each sample (x axis), we show the best fit per generation mutation rate (y axis) for varying HSC population size estimates.

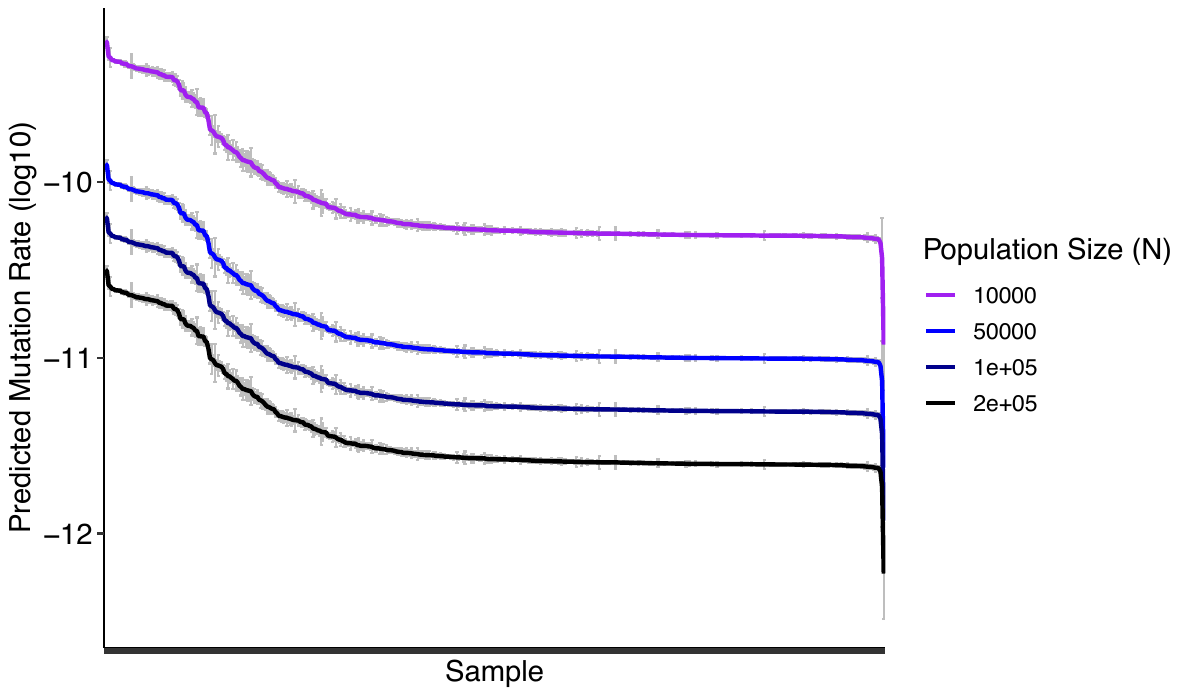

**Supplementary Figure 9. Distribution of mutations in known driver genes across evolutionary classes.** Here, we show the frequency at which driver genes harbor mutations across individuals fitting different evolutionary classes. Driver genes are shown along the x axis. Each gene is partitioned according to the frequency at which it is mutated by class (Combination: orange, Neutral: blue, Positive: green). No individuals harbouring mutations in known driver genes were predicted to be under negative selection.

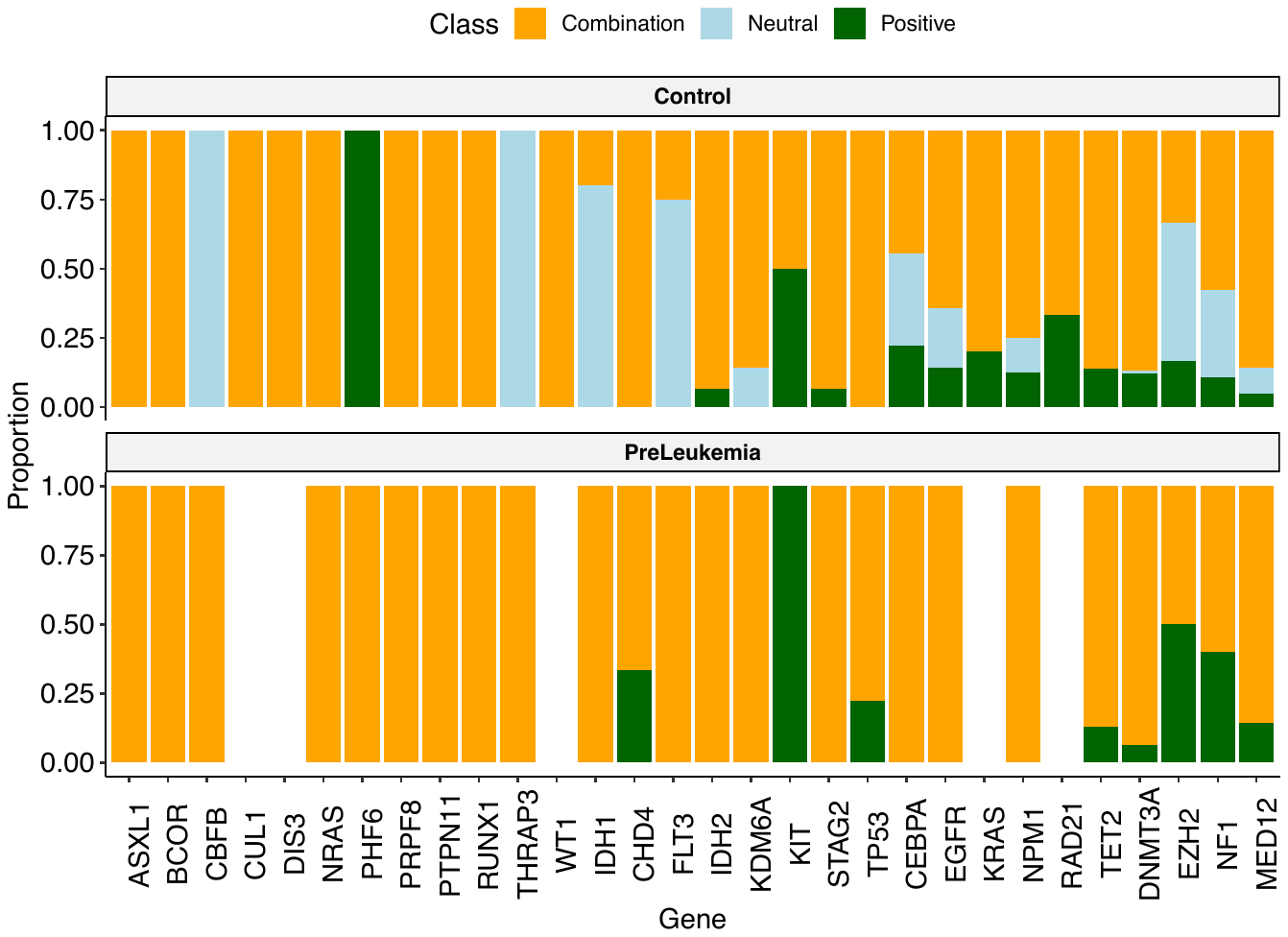
